# Fibulin-4 is highly expressed in metastatic breast cancer and can serve as a target of peptide-based imaging probes and experimental therapeutics

**DOI:** 10.1101/2025.06.18.657655

**Authors:** Alexes C. Daquinag, Solmaz AghaAmiri, Stephen M. Farmer, Sheng Zhang, Sukhen C. Ghosh, Servando Hernandez Vargas, Ashwin K. Ramesh, Zhiqiang An, Ali Azhdarinia, Mikhail G. Kolonin

**Author notes:** To whom correspondence should be addressed: Mikhail Kolonin, Ph.D. The Brown Foundation Institute of Molecular Medicine University of Texas Health Science Center at Houston 1825 Pressler St. 430E, Houston, TX 77030.

## Abstract

We have previously reported a cyclic peptide CRAGVGRGC (termed BLMP6) that homes to disseminating tumor cells in mouse cancer models and could be used for metastasis detection and intervention. Here, based on BLMP6 similarity to latent transforming growth factor beta binding protein 4 (LTBP4), we discovered fibulin-4 as a BLMP6 target. We show that BLMP6 mimics the LTBP4 domain binding to fibulin-4 and selectively binds to fibulin-4 in vitro. Fibulin-4 knockout in mouse 4T1 cancer cells abrogated BLMP6 homing to lung metastases. Fibulin-4 expression was found to be increased in invasive and metastatic human breast cancer. AZDye555 fluorophore-labeled BLMP6 was developed as a reagent selectively binding to invasive and metastatic human breast cancer cells in tissue sections and homing to MDA-MB-231 metastases in mice. We show that radiolabeling BLMP6 with ^68^Ga can be used for the detection of MDA-MB-231 metastases. We designed a peptide-drug conjugate consisting of monomethyl auristatin E (MMAE) and BLMP6 that preferentially kills aggressive cancer cells. Cytotoxicity of MMAE-BLMP6 against MDA-MB-231 tumors was confirmed *in vivo*. In an immunocompetent mouse model of B16F10 experimental lung metastases, treatment with MMAE-BLMP6 suppressed metastasis growth and improved survival. There was also a trend for metastasis suppression and survival improvement in the MDA-MB-231 experimental metastasis model. Our results suggest that fibulin-4 and BLMP6 may be further developed for the detection and targeting of metastatic human cancers.

**Statement of significance:** This study identifies fibulin-4 as a protein highly expressed in breast cancer metastasis. It evaluates the application of peptide conjugates targeting fibulin-4 in mouse models as non-invasive probes for metastasis detection and cytotoxic drug delivery.

## Introduction

Progression to metastases remains the overriding cause of mortality in patients with breast cancer (1). Metastatic cancer is not amenable to surgery and its treatment is further complicated by the development of therapy resistance often observed at advanced cancer stages (2). Early detection of metastases is therefore critical but has been limited by the lack of probes that can effectively localize them. Similar challenges persist with therapeutics selective for metastatic cancer cells. Thus, agents that specifically target pre-metastatic and disseminated tumor cells at an early stage could produce new theranostic applications and be transformative for patients with advanced breast cancer (3).

At present, detection of metastasis primarily relies on computed tomography (CT), positron emission tomography (PET), magnetic resonance imaging (MRI), X-ray, blood tests for metastasis markers, ultrasound, or biopsy analyses depending on the cancer type. Often metastases are detected too late and most patients eventually succumb to the disease due to the lack of therapeutics that can specifically target disseminated tumors. Therefore new strategies to help in early detection and real-time guidance in resection of aggressive tumors may provide improved survival outcomes for patients with advanced cancer (4). Treating metastasis is challenging due to tumor heterogeneity and cancer evolution to therapy resistance. Surgery, proton therapy, immunotherapy, chemotherapy, radiation, or immunotherapy often incompletely eradicate metastatic cells, which leads to transformation to progressively aggressive variants. Accordingly, there is significant interest in designing improved targeted imaging probes and therapeutic modalities to efficiently diagnose and treat cancer metastasis (5).

By screening a combinatorial library, we previously identified cyclic peptides that bind metastatic cells in mouse models of cancer (6). One of these peptides, CRAGVGRGC (termed BLMP6) was found to home to pre-metastatic and metastatic lesions. Colocalization with a mesenchymal N-cadherin indicated that BLMP6 is selective for cancer cells that have undergone the epithelial-to-mesenchymal transition (EMT), a process linked with metastatic dissemination. To assess the potential for *in vivo* imaging, BLMP6 was labeled with gallium (Ga) radioisotopes to quantitatively measure tissue distribution *ex vivo*. Tissue analysis showed selective uptake of ^67^Ga-BLMP6 in experimental lung metastases formed by intravenously injected cancer cells. PET imaging with ^68^Ga-labeled BLMP6 confirmed *ex vivo* findings and demonstrated the feasibility of visualizing metastases with a clinically used non-invasive modality. To test the utility of BLMP6 for delivering therapeutic payloads, we first developed a cytotoxic conjugate containing an apoptosis-inducing moiety. In pilot experiments, this hunter-killer (HK) peptide displayed cytotoxicity in cell culture and the highly aggressive mouse orthotopic graft model of 4T1 triple-negative breast cancer cells (6). While HK peptides have shown a limited therapeutic window in preclinical studies and clinical trials (7), monomethyl auristatin E (MMAE), a small molecule microtubule inhibitor, is a highly potent payload used in several FDA-approved antibody-drug conjugates (8). Therefore, MMAE-peptide conjugates may exhibit a more potent antitumor effect and have a higher potential for clinical translation.

Here, we identified fibulin-4 (FBLN4), also known as EFEMP2, as the protein target for BLMP6 binding and show that FBLN4 is highly expressed in invasive and metastatic human breast cancer. We designed a BLMP6-based probe radiolabeled with ^68^Ga for metastasis detection in a cancer xenograft model based on human MDA-MB-231 triple-negative breast cancer cells. We designed a peptide-drug conjugate for targeted delivery of MMAE to metastatic tumors. Cytotoxicity of MMAE-BLMP6 was confirmed in MDA-MB-231 cell culture and in mice. In experimental lung metastasis models, treatment with MMAE-BLMP6 also showed the potential to extend animal survival.

## Materials and Methods

### Molecular Modeling

The structures of human FBLN4 (UniProt: O95967), LTBP4-L (UniProtKB: Q8N2S1), mouse Fbln4 (UniProtKB: Q9WVJ9), Ltbp4-L (UniProtKB: Q8K4G1), and the reverse BLMP6 (CGRGVGARC) were generated by AlphaFold3 (9). PyMOL and TM-align were used to estimate the root mean square deviation (RMSD) and the template-modeling (TM)-score, respectively, measures for how similar two 3D structures are after optimal alignment based on the average distance between atoms. BLMP6 was found to be structurally similar to the 725-730 segment of mouse LTBP4 (RSMD=1.085 Å, TM-score=0.46), and the 682-687 segment of human LTBP4 (RMSD=1.860 Å, TM-score=0.46). FBLN4-BLMP6 was predicted by AlphaFold3 and CB-Dock (10), and found to have similar folding between mouse and human (RMSD=1.33 Å; TM-score=0.79). LTBP4-FBLN4 was predicted by AlphaFold3 and ClusPro (11) using a 202 amino acid fragment of FBLN4 (residues 242-443 for human and mouse) together with a 106 amino acid fragment of LTBP4 (residues 632-737 for human and 675-780 for mouse), and was found to have similar folding between mouse and human (RMSD=1.25 Å, TM-score=0.64). All models were rendered using PyMOL.

### Tissue and cell analysis

Formalin-fixed paraffin-embedded (FFPE) tissue arrays BR1008b and BR1008c were purchased from TissueArray LLC. Immunofluorescence (IF) on cells and formalin-fixed paraffin-embedded tissue sections was performed as described (12). Primary antibodies used: rabbit anti-fibulin-4 antibody EPR684(2) from Abcam (1:100), rabbit anti-Fd bacteriophage from Sigma-Aldrich (1:500), rabbit anti-Asp175-cleaved Caspase 3 from Cell Signaling Technology (1:100), rabbit anti-N Cadherin 22018-1-AP from Proteintech (1:500), mouse anti-endomucin from R&D Systems, AF4666 (1:100). Biotinylated isolectin B4 (Vector B-1205) was used at 1:50 and detected with streptavidin-Alexa Fluor 488 (Life Technologies S32354, 1:200). Secondary antibodies used: donkey Alexa Fluor 488-conjugated IgG (Invitrogen, 1:300) and Cy3-conjugated IgG (Jackson ImmunoResearch, 1:300). Terminal deoxynucleotidyl transferase TdT mediated dUTP nick-end labeling (TUNEL) was performed with a kit from ABP Biosciences (A050). Nuclei were stained with Hoechst 33258 (Invitrogen, 1:2000). Tissue and cell images were acquired with a Carl Zeiss upright Apotome Axio Imager Z1 microscope.

### Cell lines and primary cell culture

Cells were cultured in DMEM media (Hyclone) supplemented with penicillin-streptomycin (Gibco) and 10% fetal bovine serum (Atlas Biologicals). 4T1, B16F10, and MDA-MB-231 were from ATCC. 4T1 transduced to express luciferase (luc) were previously described (6). B16F10-luc-GFP and MDA-MB-231-luc-GFP cells were made by transducing parental cells with lentivirus expressing luciferase and GFP (Addgene #169308). All cells were tested for mycoplasma before expansion for experiments. For *FBLN4* knockout in 4T1 cells, a *pLenti-CRISPR/Cas9 FBLN4* gRNA vector, with 20-bp target sequence 5′-AGCGTCCCCACAGGATCCCG<u>AGG</u>-3′ containing the PAM sequence (underlined) was subcloned into *lenti-CRISPR v2* plasmid (Addgene no. 52961) and used to generate the lentivirus to knockout *FBLN4* in 4T1 cells. Primers used for measuring *FBLN4* expression: CACGGAATGCACAGATGGCTA, CATCCACACAGCTCTCCTGTT. Cell viability was measured using Celltiter-Blue assay (Promega; G8080).

### Peptide modifications

BLMP6 variants were synthesized by Genemed Synthesis Inc (San Antonio, TX) as cysteine-cyclized peptides. Resin-bound BLMP6 was synthesized in bulk to permit different bioconjugation reactions to be performed. Biotinylated BLMP6 was produced via N-terminal conjugation as previously described (6). AZDye555-BLMP6 was synthesized by conjugating AZDye555 NHS-ester dye (Vector labs; FP-1166) to the N-terminus of BLMP6 in anhydrous DMF in the presence of diisopropyl amine at 37°C for 3 h, followed by overnight stirring at room temperature. The conjugate was purified by reversed-phase high-performance liquid chromatography (RP-HPLC) with a mobile phase of water and acetonitrile containing 0.1% trifluoracetic acid. Fractions containing the desired product were collected and lyophilized to yield purified AZDye555-BLMP6. Product identity was confirmed by mass spectra analysis: ESI-MS (+) *m*/*z*: calcd 1691.07; found 1692.19 (M + 2H)^2+^. To produce the radioactive compounds, the chelating agent 1,4,7,10-tetraazacyclotetradecane-*N′,N″,N‴,N″″*-tetra-acetic acid (DOTA) t-Bu ester was conjugated to resin-bound BLMP6 and labeled with ^68^GaCl_3_ as described (6). For MMAE-BLMP6 synthesis, an all-D-amino acid analog of BLMP6 was synthesized. MMAE was coupled to a linker PAB-Cit-Val-PEG4-NHS consisting of PEG4, valine-citrulline, and p-aminobenzyl (PAB), ordered from Creative Biolabs (Shirley, NY). For MMAE conjugation, BLMP6 (0.004 mmol) was dissolved in DMF (1 mL) mixed with diisopropylethylamine (5 µL), and the cytotoxic payload moiety MMAE-PAB-Cit-Val-PEG4-NHS (0.003 mmol). The reaction mixture was stirred for 4 h at 37°C and monitored by analytical HPLC. After purification, compound identity was validated by mass spectra analysis: ESI-MS (+) *m*/*z*: calcd 2260.8; found 1129.58 (M + 2H)^2+^.

### Peptide binding assays

Octet binding assays were performed with an Octet RED96 system (ForteBio) as described (13). Briefly, the peptides were loaded onto streptavidin resin-coated biosensors (Sartorius Lab, 18-5019) at a concentration of 20 ug/ml in 1x kinetic buffer for 5 min. The coated biosensors were then incubated with a series of Fibulin-4 protein (Abcam, 182808) concentrations (ranging from 0 to 400 nM) for 300 sec, followed by a washing step in 1x kinetic buffer for an additional 300 sec to allow for dissociation. For all experiments, Ka (1/Ms) and Kd (1/s) were measured. FortéBio’s data analysis software was employed to fit the binding curve to a 1:1 binding model in order to extract association and dissociation rates. Overlay assay on tissues was performed by incubating AZDye555-BLMP6 (12.5 ug/ml) for 2 hr and washing with PBS before fixation with 4% paraformaldehyde.

### Animal experiments

Animal studies were approved by the IACUC of UTHealth. Spontaneous cancer progression to metastases was modeled by injecting 2×10^4^ 4T1 cells into the mammary fat pad of 10-week-old female Balb/c mice (Jackson Laboratory) with a 31-gauge needle. Metastatic progression was monitored with bioluminescence imaging. For lung metastasis modeling, Balb/c mice were tail vein-injected with 2.5×10^5^ 4T1-luc WT or FBLN4-KO cells, C57BL/6 mice (Jackson) were tail vein-injected with 1×10^5^ B16F10 cells, and athymic Nude-Foxn1nu (Envigo) mice were tail vein-injected with 1×10^5^ MDA-MB-231-luc-GFP cells. For phage-peptide homing analysis, mice were injected into the tail vein with 5×10^10^ transducing units (TU) of phage-BLMP6 (6). After 30 min of circulation, heart perfusion with 10 ml of PBS was performed and lung sections were processed. For PET imaging studies, mice were intravenously injected with 7.4 MBq (200 µCi, 5 nmol) of ^68^Ga-labeled DOTA-BLMP6 and imaged 45-60 min later using Albira small-animal PET/CT scanner (Bruker After imaging, the mice were euthanized, the tissues of interest were harvested and probe homing was analyzed using a Cobra II automated gamma counter (Packard) and expressed as percentage of the injected dose per gram tissue (%ID/g tissue).

### Statistical Analysis

GraphPad Prism v.9.0.2 was used to graph data as mean ± S.E.M. Statistical significance was determined with unpaired t-test or two-way ANOVA built-in analysis tools. P-values (*p*) <0.05 were considered significant.

## Results

### BLMP6 mimics LTBP4 and binds to Fibulin-4

Analysis of mammalian proteins for similarity for BLMP6 by BLASTP identified mouse latent transforming growth factor beta binding protein 4 (LTBP4). Amino acids 725-730 within the EGF-like calcium-binding domain 6 of LTBP4 matched the reverse BLMP6 sequence CGRGV. Molecular modeling by AlphaFold3 predicted that BLMP6 is structurally similar to this 725-730 segment of mouse LTBP4 (Figure 1A, RSMD=1.085 Å, TM-score=0.46), and the conserved 682-687 segment of human LTBP4, CGRGA (Figure S1A, RMSD=1.860 Å, TM-score=0.46). LTBP4 is known to bind to the carboxy (C) terminus of fibulins (14). Specifically, FBLN4 has been reported to interact with the EGF-like calcium-binding domain 6 of LTBP4 (15), which corresponds to the region of BLMP6 mapping. Using AlphaFold3 and CB-dock, we simulated this binding (Figure 1B and S1B) and found that BLMP6 occupies a similar site on FBLN4 in mouse and human (RMSD=1.33 Å; TM-score=0.79) by engaging with R326, R346, I363, Q364, T366, and S367. Both AlphaFold3 and ClusPro confirmed that LTBP4 engages with this same region on FBLN4 (Figure 1B and S1B), with a similar pose predicted for both mouse and human (RMSD=1.25 Å, TM-score=0.64).

**Figure 1.**
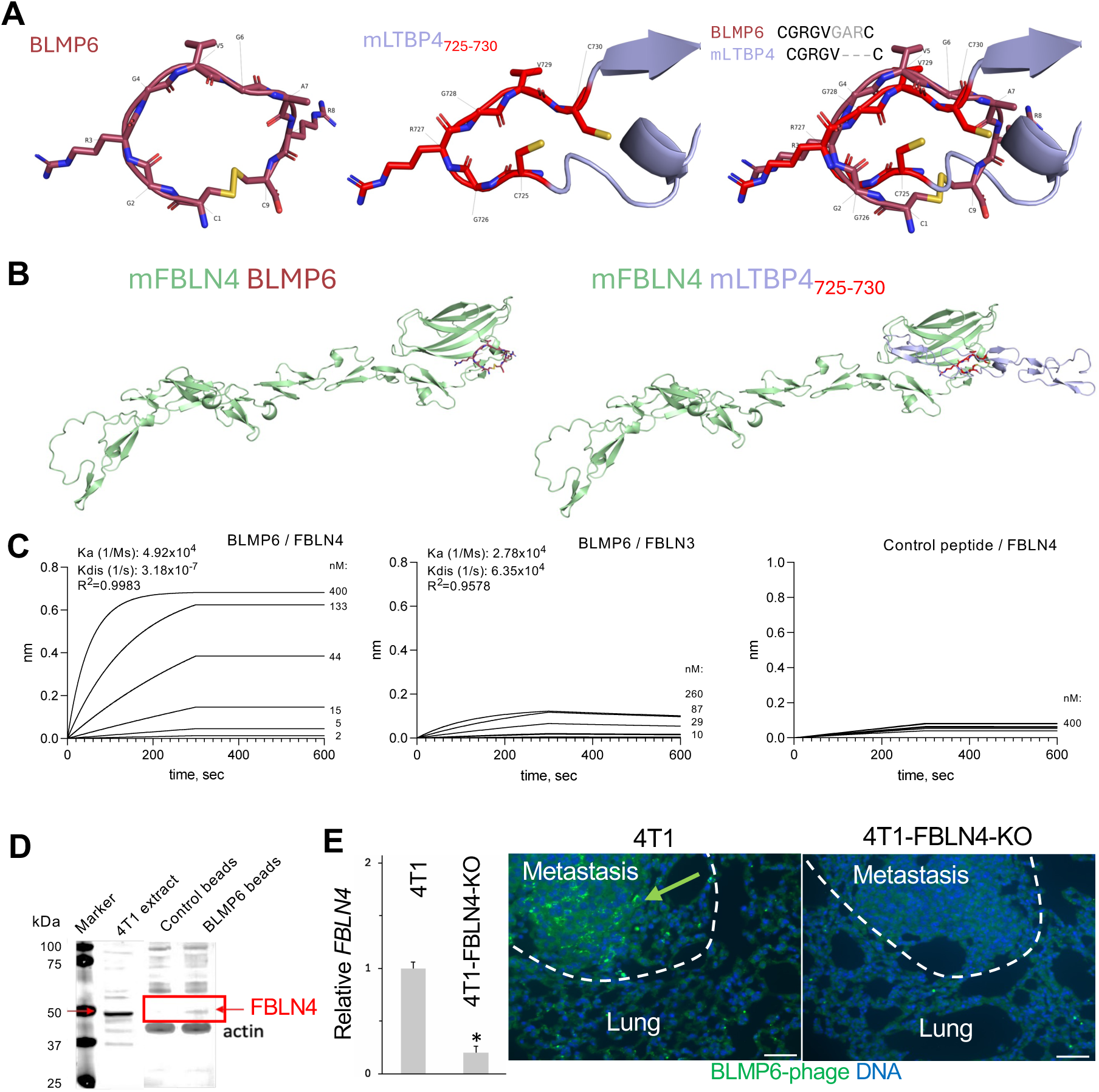
BLMP6 mimics LTBP4 and binds to FBLN4. **A**, AlphaFold3 (AF3) modeling of the 3D structure of BLMP6 and mouse LTBP4L (residues 716-733 shown) containing the BLMP6 similarity segment (725-730, highlighted). **B**, Left: consensus docking pose predicted by AF3 and CB-Dock shows that BLMP6 interacts with the C-terminus of mouse FBLN4 (residues 41-443 shown). Right: a similar binding predicted for a 106 amino acid fragment of LTBP4L (residues 675-780) containing the BLMP6 similarity segment (725-730, highlighted) using AF3 and ClusPro. **C**, Biolayer interferometry assay performed with Octet to measure binding affinities. **D**, Affinity purification of FBLN4 from 4T1 cell extract with biotinylated BLMP6 immobilized on streptavidin-coated beads. Immunoblotting with anti-FBLN4 antibodies identifies the expected 49kDa protein, which is not isolated on control beads. **E**, Graph: RT-PCR analysis of gene mRNA expression, relative to *18S*, confirming the knockout of FBLN4 in 4T1 cells. Images: lung paraffin sections from mice that were injected with phage displaying BLMP6. Anti-phage IF demonstrates BLMP6-phage homing to 4T1 metastases but not to 4T1-FBLN4-KO metastases FBLN4-KO metastases. Scale bar: 100 μm.

To test if BLMP6 binds to FBLN4 *in vitro*, we used Octet to perform the biolayer interferometry assay. The binding affinity of BLMP6 binding to FBLN4 was found to be three orders of magnitude higher than binding to FBLN3 (Figure 1C). In contrast, a cyclic peptide CKGGRAKDC (16) used as a control, displayed no binding to FBLN4 (Figure 1C). To confirm BLMP6 binding to FBLN4 expressed by cancer cells, proteins extracted from 4T1 cells were incubated with beads pre-loaded with BLMP6. Upon washing and elution, the predicted 49 kDa band was observed by immunoblotting with anti-FBLN4 antibodies (Figure 1D). To confirm BLMP6 binding to FBLN4 in a genetic model, we used CRISPR/Cas9 to make 4T1 FBLN4 knockout (KO) cells confirmed by RT-PCR (Figure 1E). FBLN4-KO and control 4T1-luc cells were grafted into mice, and once metastases were detected by bioluminescence, phage displaying BLMP6 (6) was injected. After 30 min of circulation, heart perfusion was performed, and lungs were recovered. Section histopathology confirmed metastasis formation in all lungs, indicating FBLN4 expression is not required for cancer cell colonization and growth at the secondary site. Immunofluorescence (IF) analysis with an anti-phage antibody confirmed BLMP6-phage homing to 4T1 metastases, whereas no BLMP6-phage was observed in FBLN4-KO metastases (Figure 1E). These data indicate that FBLN4 is the receptor of BLMP6.

### AZDye555-BLMP6 as a reagent to identify invasive breast cancer in tissue sections

In our previous studies, cyclic peptides conjugated with fluorophores have been used to identify cells of interest in paraffin-embedded tissue sections (17). We next tested if BLMP6 labeled with a red AZDye555 fluorophore, can be used as an antibody to identify invasive cancer cells. AZDye555-BLMP6 was confirmed to bind to HEK293 cells ectopically expressing FBLN4 but not to non-transfected control cells (Figure S2A). When applied to lung sections from mice grafted with mouse 4T1 cells, AZDye555-BLMP6 selectively bound to metastatic lesions but not to normal lung (Figure 2A). The signal was observed exclusively on endomucin-negative cells, indicating that BLMP6 does not bind to the vasculature. We previously reported that biotinylated BLMP6 binds human triple-negative breast cancer (TNBC) MDA-MB-231 cells (6). To test if AZDye555-BLMP6 can be used to identify human cancer *in situ*, we applied it to lung sections from mice pre-injected with MDA-MB-231 and growing experimental lung metastases. Lung section IF demonstrated that AZDye555-BLMP6 binds to a subpopulation of tumor cells expressing N-cadherin, a marker of the EMT (Figure 2B). Notably, the non-malignant stroma, also expressing N-cadherin, was devoid of AZDye555-BLMP6 binding (Figure 2B). We also tested this method on human tissue arrays containing 50 primary and 40 metastatic breast cancer biopsies as well as 10 adjacent control breast tissue biopsies (Figure S2B-C). IF analysis with N-cadherin antibodies detected its expression in epithelial cells, which was at a background level in normal breast tissue and low-grade tumors but induced in most stage II-III tumors, as well as metastases (Figure 2C). Notably, AZDye555-BLMP6 binding was largely concordant with N-cadherin expression observed across the cores (Figure 2C and S2C). In invasive stage III tumors and metastasis, AZDye555-BLMP6 binding was particularly prominent and co-localized mainly with malignant epithelial cells, whereas some of the stromal cells expressing N-cadherin were devoid of AZDye555-BLMP6 binding (Figure 2C). These data are consistent with BLMP6 binding to invasive and metastatic human cancer cells that have undergone the EMT.

**Figure 2.**
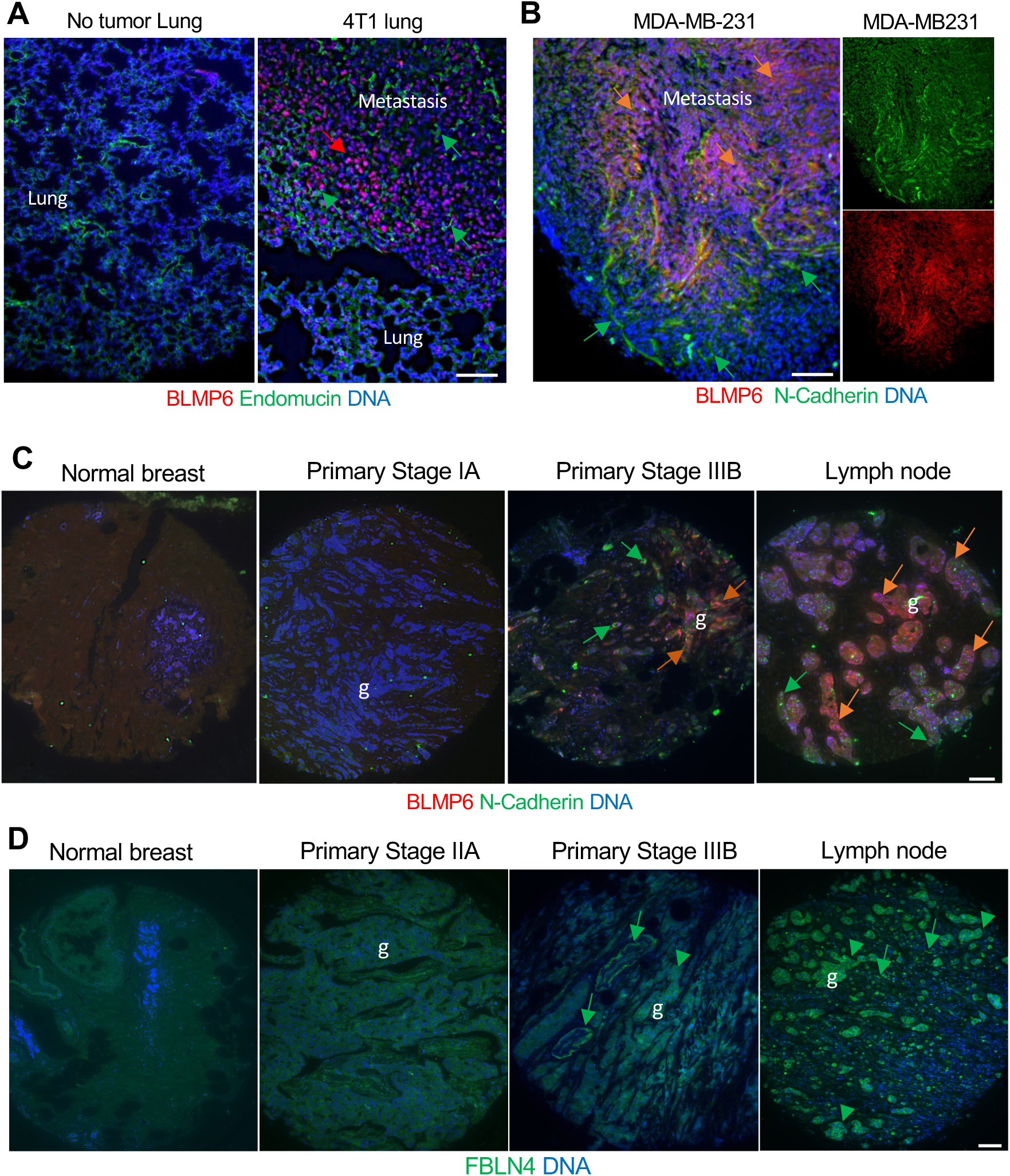
BLMP6 binds to invasive breast cancer cells in tissue sections. **A**, Lung paraffin sections from mice without or with 4T1 metastases overlayed with AZDye555-BLMP6 (12.5 ug/ml) reveal binding specificity to metastatic cells. Endomucin IF identifies endothelial cells. **B**, Lung paraffin section from a mouse pre-injected (tail vein) with MDA-MB-231 cells overlayed with AZDye555-BLMP6 (12.5 ug/ml) reveals binding to metastatic cells. N-cadherin IF identifies AZDye555-BLMP6-bound cancer cells that underwent EMT (orange arrows) and stromal cells (green arrows). **C**, Representative cores from human FFPE tissue array BR1008b overlayed with AZDye555-BLMP6. N-cadherin IF identifies AZDye555-BLMP6-bound glandular cancer cells (g) that underwent EMT (orange arrows) and stromal cells (green arrows). **D**, Representative cores from human FFPE tissue array BR1008c. IF identifies FBLN4 in glandular cancer cells that underwent EMT (arrowheads) and in the stroma (arrows). Scale bar: 500 μm.

FBLN4 expression in cancer cells being a requisite for it serving as a target for BLMP6 metastasis homing suggested that cancer progression is linked with changes in FBLN4 expression. We investigated this by using the human tissue arrays (Figure S2C). IF analysis with anti-FBLN4 antibodies detected its expression in epithelial cells, which was low in normal breast tissue and generally low in the majority of tumor samples from stage II tumors (Figure 2D). In contrast, FBLN4 expression was found to be higher in the epithelium and present in the stroma, of many invasive stage III tumors (Figure S2C). The majority of metastasis biopsies displayed high FBLN4 expression in the epithelium and the stroma (Figure 2D). Figure S2D summarizes BLMP6 binding and FBLN4 expression in the 100 tissue biopsies. These data indicate that FBLN4 expression, induced during cancer progression, is consistent with the selectivity of BLMP6 for invasive cancer cells.

### Detection of human metastases in mice with BLMP6 probes

In our previous study, BLMP6 labeled with a Cy3 fluorophore was shown to home to experimental mouse lung metastases (6). To test if BLMP6 can home to metastases of human cells, we used our newly developed AZDye555-BLMP6 reagent. Specifically, after mice tail vein-injected with MDA-MB-231 cells expressing GFP and luciferase were confirmed to develop lung metastases by IVIS, intravenous injection of AZDye555-BLMP6 was performed. Sections of lung tissues confirmed GFP-positive metastatic lesions (Figure 3A). AZDye555-BLMP6 signal was detected within the lesions but not in the surrounding normal lung, confirming BLMP6 homing to human metastases in mice.

**Figure 3.**
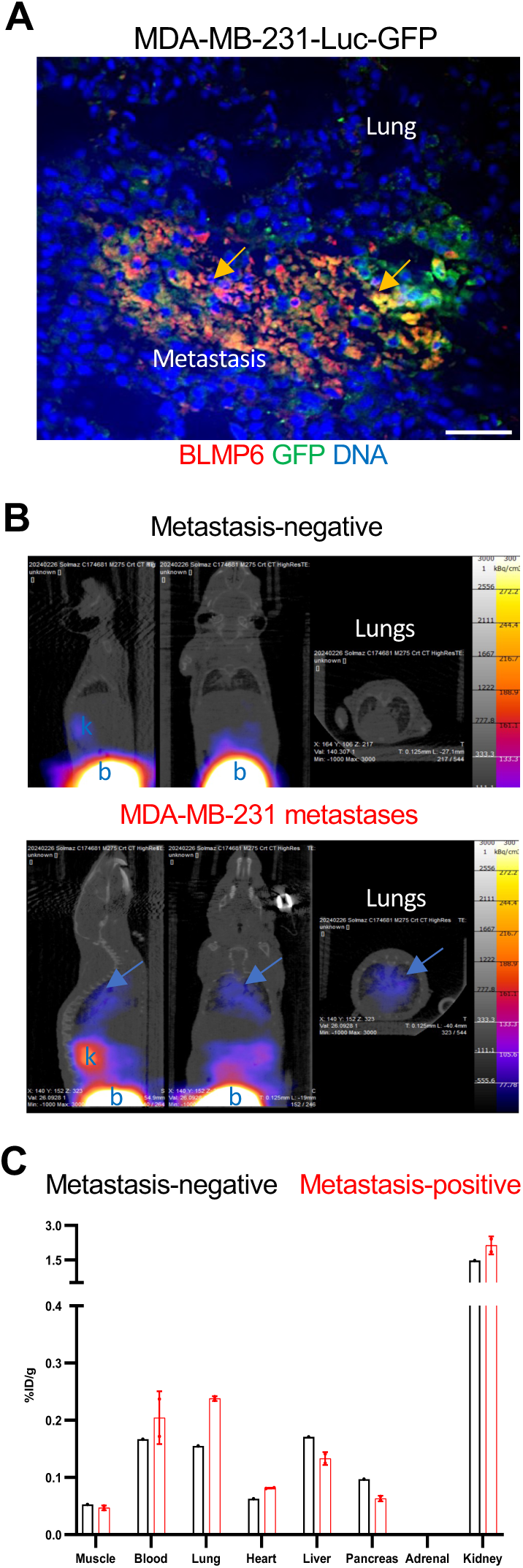
BLMP6 homes to human cancer metastases. As an experimental model, immunodeficient mice intravenously (iv)-injected with MDA-MB-231-Luc-GFP cells were used upon lung metastasis detection by IVIS. **A**, A lung section from a mouse iv-injected with AZDye555-BLMP6 (50 ug) shows a GFP-positive metastatic lesion with AZDye555-BLMP6 signal, absent in the surrounding non-tumor tissue. **B**, Radiolabeled ^68^Ga-BLMP6 iv-injected into nude mice without and with MDA-MB-231 lung metastases detected by IVIS. At 1 h, PET/CT images are scaled equally (0.4-1.96 %ID/g) to allow cross-reference throughout the study. Arrow: ^68^Ga-BLMP6 in metastases. b: bladder, k: kidney **C**, Biodistribution in tissues resected from mice without and with metastases (B) with gamma counter plotted as percentage of the injected dose per gram tissue (%ID/g tissue).

In our previous study, Ga-labeled BLMP6 was also used to detect BLMP6 metastasis homing by PET/CT and to quantitatively measure tissue distribution (6). We first used the experimental lung metastasis model of B16F10 melanoma cells injected intravenously to standardize the protocol for radiolabeled peptide probes. BLMP6 was conjugated with the radiometal chelator DOTA, which was then used for radiolabeling. As expected, a high ^68^Ga-BLMP6 signal was observed in the bladder and the kidneys excreting the radiolabeled peptides; the signal in most other organs was similar to the background, indicating low non-specific binding (Figure S3A). ^68^Ga-BLMP6 lung signal above the background level was observed in mice with B16F10 lung metastases but not in tumor-free mice (Figure S3A). *Ex vivo* tissue quantification on a gamma counter confirmed the imaging findings (Figure S3B). We then used an orthotopic mouse model of TNBC to test BLMP6 homing to spontaneous pulmonary metastases. Three weeks after grafting 4T1 cells expressing luciferase (luc) into the mammary pad of Balb/c mice, we performed the experiments with ^68^Ga-BLMP6, as for the B16F10 model. Again, bladder and kidney signals were comparable for mice with and without metastases (Figure S3C-D). ^68^Ga-BLMP6 signal was detected in the lungs only when metastases were observed (Figure S3C). ^68^Ga-BLMP6 was also detected in the primary tumor (Figure S3C-D), which complicated imaging. Therefore, for testing BLMP6 as a probe in the lung metastasis model of human TNBC, MDA-MB-231-Luc-GFP cells were injected intravenously into athymic mice. When metastases became detectable by IVIS, we performed the experiments with ^68^Ga-BLMP6 as above. As for allograft models, bladder, kidney, and control organ signals were at the background levels in tumor-positive and control mice (Figure 3B-C). Accumulation of ^68^Ga-BLMP6 above the background level was detected in the lungs of mice with MDA-MB-231 metastases but not in tumor-free mice (Figure 3B-C). These data indicate that BLMP6 homes to metastases of human cells.

### MMAE-BLMP6 targets and suppresses metastases

We designed a BLMP6-drug conjugate using the same components as validated MMAE-based antibody-drug conjugates (18–22). MMAE was coupled to BLMP6 *via* a linker consisting of PEG4 (added hydrophilicity), valine-citrulline (cathepsin-sensitive cleavable segment), and PAB (self-immolating segment) as described (19–25). This well-characterized cleavable linker strategy achieves intracellular drug release and favors renal elimination, hence reducing cytotoxicity in normal tissues (26,27). BLMP6 synthesized with all amino acids as D-enantiomers was used to make the compound stable to proteases and inert for the immune system (28) as previously described (29–31). Analytical testing of MMAE-BLMP6 confirmed product identity and >95% chemical purity for subsequent pharmacological tests (Figure S4A). First, the efficacy of MMAE-BLMP6 was tested in cell culture. We used the quantitative CellTiter-Blue cell viability assay (25) to quantify cell death 24 h after adding MMAE-BLMP6 to the cell culture medium. The 50% inhibitory response (IC_50_) was 500 nM for 4T1 cells (Figure 4A). Lower cytotoxicity against FBLN4-KO cells (Figure 4A) confirmed that MMAE-BLMP6 uptake is mediated by FBLN4 binding. Human MDA-MB-231 cells were killed by MMAE-BLMP6 with IC_50_ of 120 nM (Figure 4B), whereas BLMP5, another TNBC-homing peptide (6) coupled to MMAE had IC_50_ of 400 nM. MMAE-BLMP6-induced cell death for each line was confirmed by microscopy upon Trypan blue staining (Figure 4A-B).

**Figure 4.**
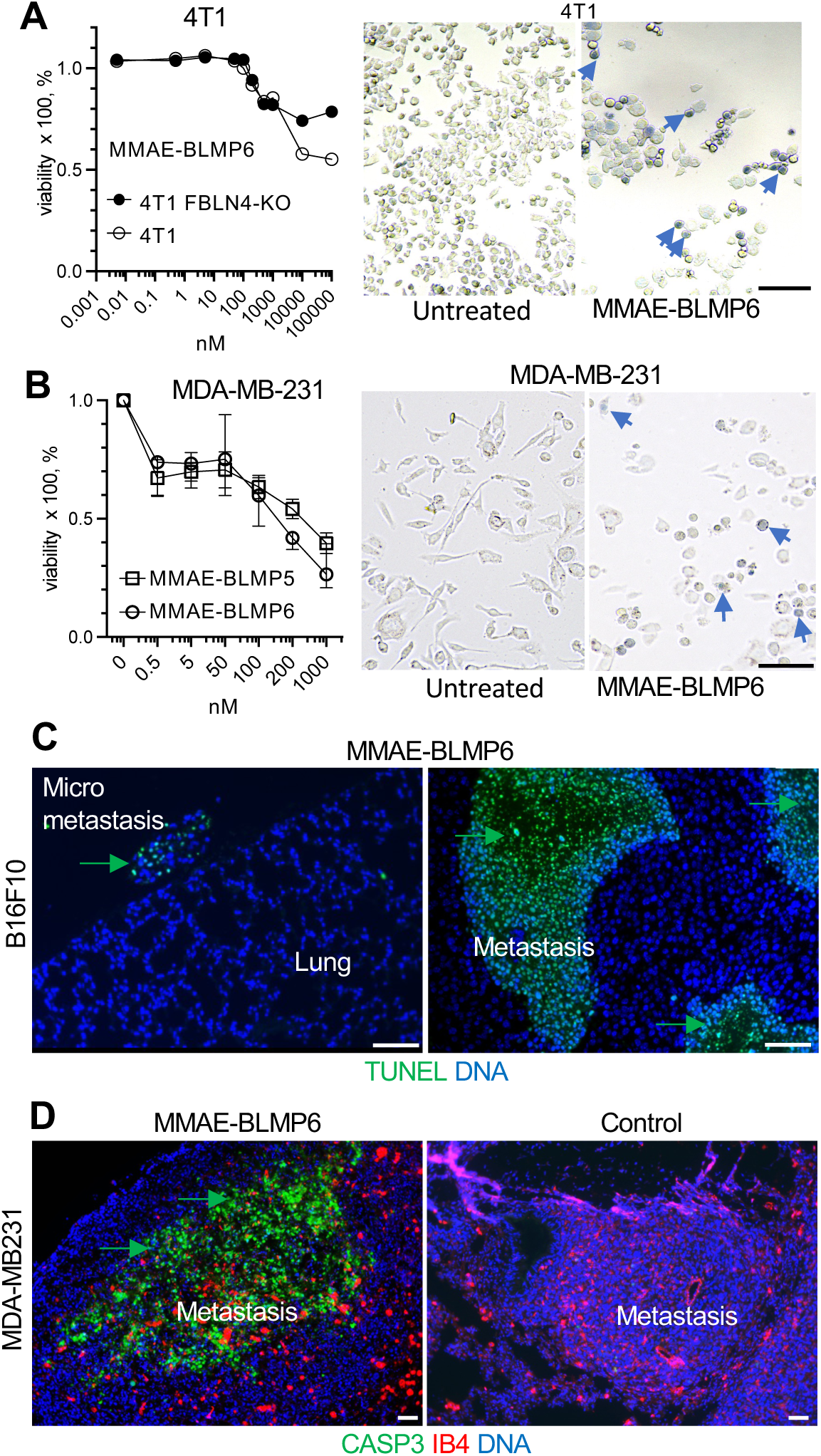
MMAE-BLMP6 targets and suppresses metastases. **A.** Survival of 4T1 and 4T1-FBLN4-KO cells treated with indicated concentrations of MMAE-BLMP6 for 24 h measured by CellTiter-Blue cell assay. Images: trypan blue staining of 4T1 cells untreated or treated with 1 μM MMAE-BLMP6. **B**, Survival of MDA-MB-231 cells treated with indicated concentrations of MMAE-BLMP6 and MMAE-BLMP5 peptides for 24 h measured by CellTiter-Blue cell assay. Images: Images: trypan blue staining of MDA-MB-231 cells untreated or treated with 1 μM MMAE-BLMP6. **C**, Lung paraffin section from mice with B16F10 metastases injected (sc) with 20 μg of MMAE-BLMP6 and sacrificed 24 hr later. TUNEL assay identifies apoptosis in metastatic cells (arrows). **D**, Lung paraffin section from mice with MDA-MB-231-luc-GFP metastases injected (sc) with MMAE-BLMP6. IF with an antibody against cleaved caspase 3 identifies apoptosis in metastatic cells (green arrows). IB4: isolectin B4 (Cy3-conjugated) binding marking the endothelium.

To test MMAE-BLMP6 efficacy *in vivo,* we first analyzed its stability in mouse serum. After 1 hr, the single MMAE-BLMP6 peak was observed, and only after 6 hrs a secondary degradation peak appeared (Figure S4B-C). We then used the experimental lung metastasis model of B16F10 cells injected intravenously to produce lung metastases. At 24 h after subcutaneous injection of MMAE-BLMP6 (20 μg), apoptosis was observed by TUNEL staining in metastatic cells (Figure 4C). In the MDA-MB-231 experimental lung metastasis model, cell death was also observed by IF detecting cleaved caspase 3 in metastases of mice treated with MMAE-BLMP6, but not in untreated mice (Figure 4D). These data indicate that MMAE-BLMP6 selectively kills metastatic cancer cells *in vivo*.

To test MMAE-BLMP6 efficacy for cancer suppression, we first used the B16F10 experimental lung metastasis model. Three MMAE-BLMP6 (50 ug) or PBS subcutaneous injections were done over days 1-6 and four additional MMAE-BLMP6 (100 ug) or PBS subcutaneous injections on days 8-18 after iv injection of B16F10-luc cells (Figure 5A). Bioluminescence analysis by IVIS revealed metastasis suppression in MMAE-BLMP6-treated mice at day 5 (Figure 5B). While vehicle-treated mice began dying on day 15, with most animals perishing by day 21, MMAE-BLMP6-treated mice only started showing metastases on day 15 by IVIS and survived 2 to 3 days longer (Figure 5A). In the MDA-MB-231-luc-GFP experimental lung metastasis model, a similar trend was observed (Figure 5C). Because mice treated with 50 to 100 μg of MMAE-BLMP6 started displaying signs of sickness at day 15, treatment was interrupted for 6 days and then re-initiated with a lower dose (15 μg daily). Bioluminescence imaging indicated that MMAE-BLMP6 treatment suppressed metastasis growth (Figure 5D). While there was no statistical difference in the overall survival curves, one of the MDA-MB-231-luc-GFP treated mice outlived the controls by 9 days (Figure 5C).

**Figure 5.**
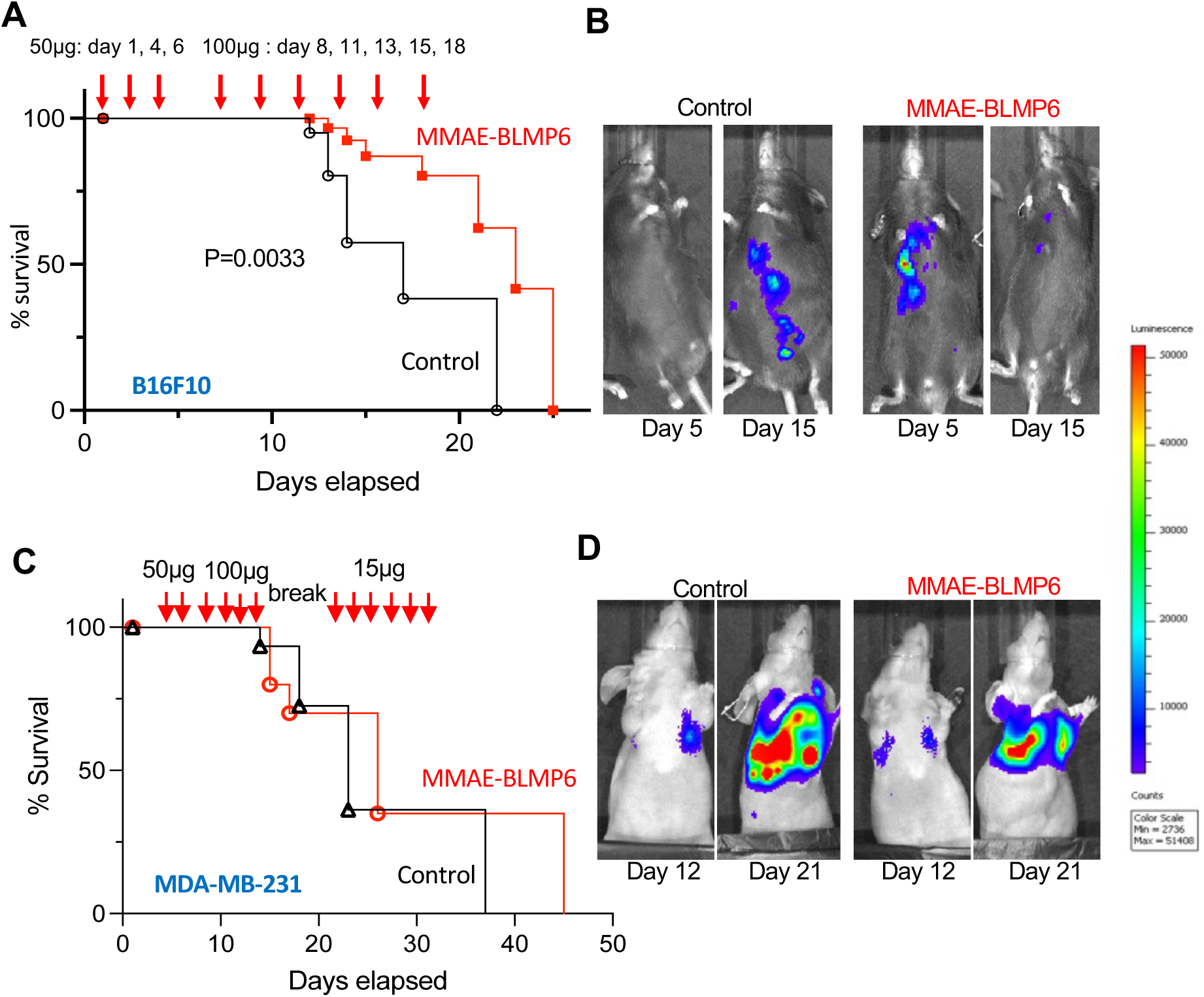
MMAE-BLMP6 suppresses metastasis progression. **A**, Kaplan-Meier curve shows the cumulative survival of mice iv-grafted with B16F10-luc cells and subcutaneously (sc)-injected with MMAE-BLMP6 or PBS as indicated. **B**, Bioluminescence analysis by IVIS revealed metastasis suppression in MMAE-BLMP6-treated mice at days 5 and 15. **C**, Kaplan-Meier curve shows the cumulative survival of mice iv-grafted with MDA-MB-231-luc-GFP cells and sc-injected with MMAE-BLMP6 or PBS as indicated. **D**, Bioluminescence analysis by IVIS revealed metastasis suppression in MMAE-BLMP6-treated mice at days 12 and 21.

## Discussion

This study was founded on BLMP6, a peptide that previously was developed for the detection and targeting of breast cancer metastases (6). Here, we show that labeled BLMP6 peptides can be used for targeting metastases in a human cancer xenograft model. We demonstrate that AZDye555-BLMP6 selectively binds to cells that have undergone EMT in invasive breast cancers and can be used to identify them not only in mice but also in paraffin sections of human tumors. We have designed a peptide-directed drug by conjugating BLMP6 to MMAE and demonstrated its tolerability at concentrations higher than those observed for untargeted MMAE (25,32). We also present evidence of efficacy in mouse metastasis models. It should be noted that since the TNBC models chosen are very aggressive the observed response is promising. The range of doses at which MMAE-BLMP6 is effective and tolerable *in vivo* is consistent with that reported for MMAE peptide and antibody conjugates (25,32). Binding was also observed in primary tumors expressing N-cadherin, stages post-EMT.

Our data indicate that BLMP6 mimics a segment of LTBP4, a large ECM glycoprotein that binds latent TGF-β complexes and localizes them to the ECM (15). LTBP4 interacts with FBLN4 via its C-terminal domain, which overlaps with binding sites for other ECM proteins. FBLN4, a member of the fibulin family of matricellular glycoproteins, has been identified as the receptor of BLMP6. Our data confirms the affinity and selectivity of BLMP6 binding to FBLN4. FBLN4 is not typically expressed on the cell surface. Instead, it is secreted into the ECM. However, it has been reported that FBLN4 can be internalized by endocytosis (33). This offers a potential mechanism for BLMP6-mediated cancer cell targeting. The interaction between LTBP4 and FBLN4 is important in the context of ECM organization, and TGF-β signaling regulation. Molecular modeling predicted that BLMP6 docks the C-terminus of FBLN4 by binding R326, R346, I363, Q364, T366, and S367 in the pocket also bound by LTBP4. It remains to be determined why BLMP6 binds to FBLN4 selectively in the context of invasive and metastatic cancer. LTBP4 has two major isoforms: the long form (LTBP4L) and the short form (LTBP4S). Both isoforms can bind to FBLN4, but LTBP4L exhibits a stronger binding affinity (34). Moreover, it has been shown that LTBP4L has higher avidity for FBLN4 multimers (14). Therefore, multimerization of FBLN4 upon its increased secretion by invasive cancer cells could account for the BLMP6 binding selectivity. FBLN4 is also known to undergo post-translational modifications (PTMs) that are crucial for its proper function, stability, and interactions with other proteins. These include alternative disulfide bond formation, N-linked glycosylation, and proteolytic processing. Matrix metalloproteinases (MMPs) have been reported to proteolytically cleave FBLN4 into fragments, with MMP-7 and MMP-12 being particularly potent (35). These PTMs could also contribute to the selectivity of FBLN4 targeting in invasive cancer.

In this study, we identify FBLN4 as a potential marker and drug target in invasive breast cancer. FBLN4 plays a key role in elastic fiber formation and the stabilization of ECM components. It has been shown to regulate the activation of lysyl oxidase, an enzyme responsible for cross-linking elastin and collagen (33). FBLN4 has emerged as an important matricellular factor that can promote or suppress cancer progression depending on the cancer type. High FBLN4 expressed correlates with advanced disease stages, poor differentiation, lymph metastasis, and unfavorable prognosis of cervical and ovarian cancer (36). It has been shown that FBLN4 promotes the EMT and that FBLN4 knockdown reduces osteosarcoma metastasis (37). Importantly, the expression of FBLN4 in breast cancer has been reported (36), including proteomic analysis that identified FBLN4 among proteins differentially secreted in metastatic breast cancer cells (38). However, its role in metastasis progression has not been investigated. Understanding the mechanisms underlying FBLN4 processing could inform further development of targeted therapies and diagnostic tools. Overall, the potential utility of BLMP6 as a vehicle for selective agent delivery to invasive cancer cells could be pursued along with FBLN4 antibodies.

## Acknowledgments

We thank Yongmei Yu, Suridh Adhikari, and Sheng Pan for the help with the experiments. We thank Charles Kingsley, Emily Newsom, Mark Cabrera, Xavier Sauceda, and Jennifer Meyer for their assistance with the PET/CT imaging studies, and the University of Texas MD Anderson Cancer Center Small Animal Imaging Research Facility personnel for help with radiolabeled peptide imaging studies.

This work was supported by DOD BCRP grant BC211016 to M.G.K. and A.A., Welch Foundation grant AU-0042-20030616 to Z.A., and CPRIT grant RP190561 to Z.A.

