## Supplemental FIgures for "Fibulin-4 is highly expressed in metastatic breast cancer and can serve as a target of peptide-based imaging probes and experimental therapeutics"

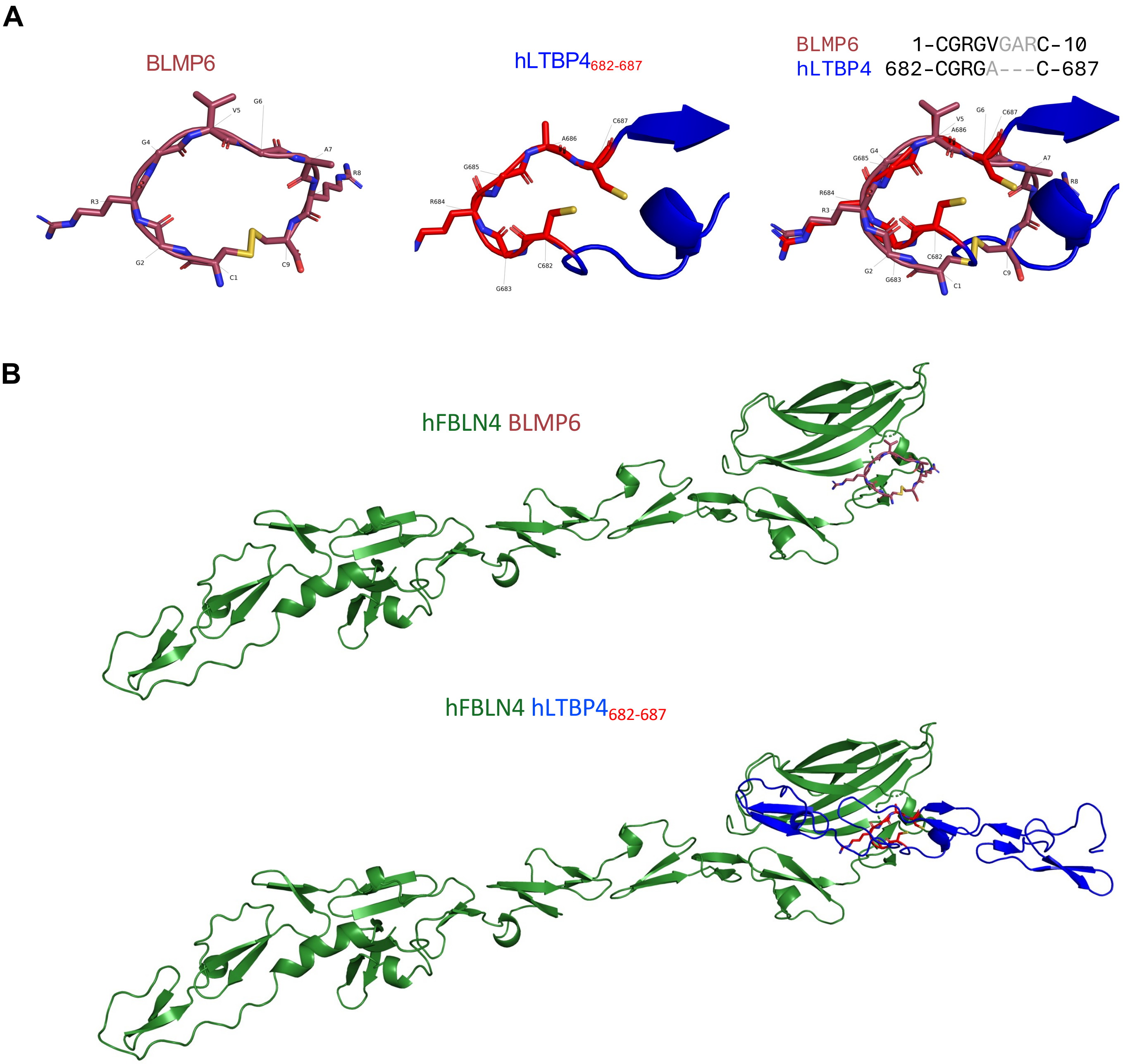


**Figure S1.** BLMP6 mimics LTBP4 and binds to FBLN4. **A**, AlphaFold3 (AF3) modeling of the 3D structure of BLMP6 and human LTBP4L (residues 673-690 shown) containing the BLMP6 similarity segment. **B**, A consensus docking pose predicted by AF3 and CB-Dock shows BLMP6 binding to the C-terminus of full-length human FBLN4 (residues 41-443 shown), revealing a similar binding site to that predicted by AF3 and ClusPro for a 106 amino acid fragment of human LTBP4L (residues 632-737).

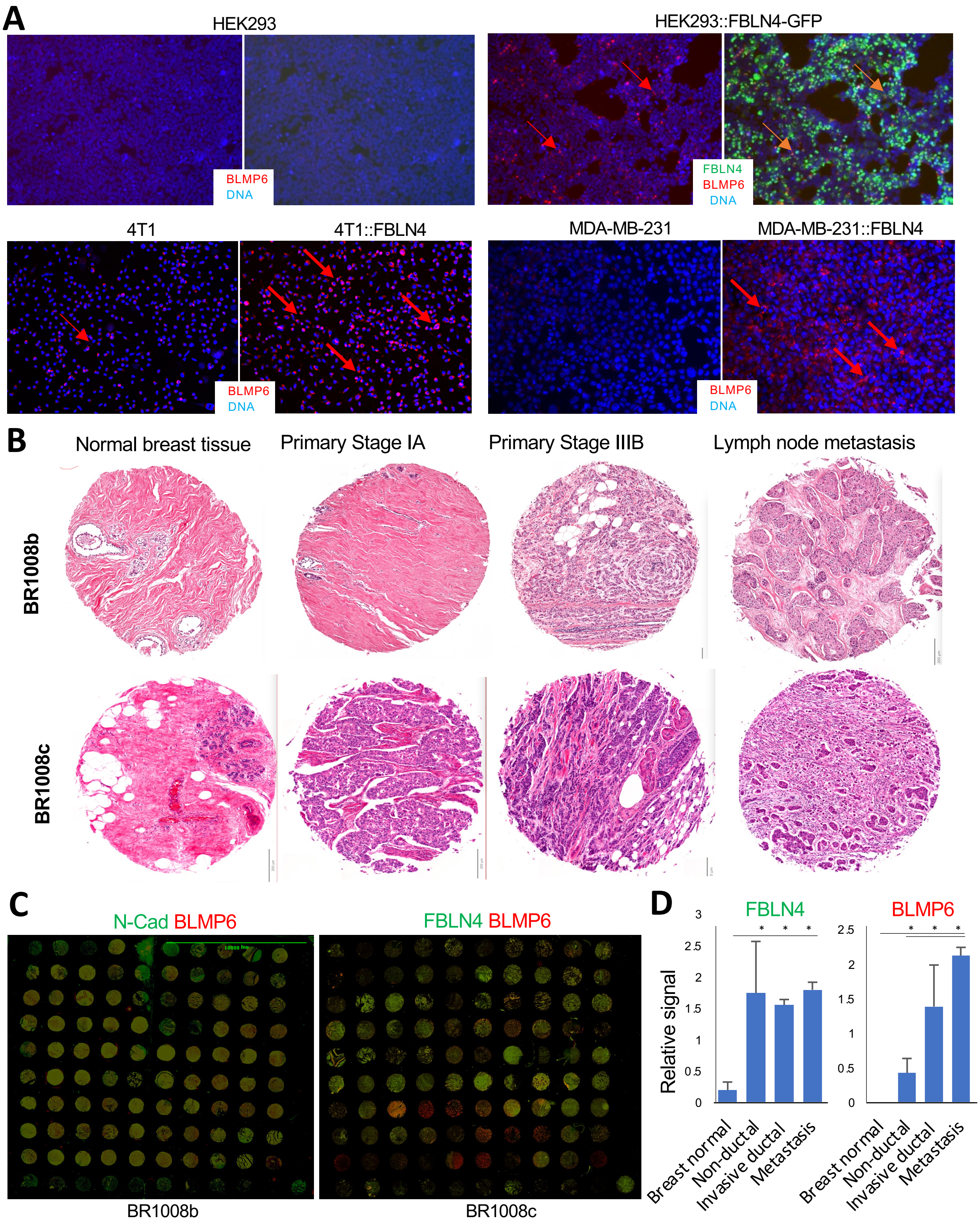


**Figure S2. A**, HEK293 cells transfected with a plasmid for expression of FBLN4-GFP fusion protein (Sino Biological; MG52071-ACG). 4T1 and MDA-MB-231 transduced with a lentivirus for expression of FBLN4 protein. AZDye555-BLMP6 was added to the growth media. After o/n incubation cells were washed with PBS, fixed, and imaged. **B**, Hematoxylin/eosin stainings of serial sections for BR1008b cores shown in Figure 2C and of BR1008c cores shown in Figure 2D. **C**, Array BR1008b analyzed for N-cadherin IF / AZDye555-BLMP6 binding. Array BR1008c analyzed for FBLN4 IF / AZDye555-BLMP6 binding. **D**, Quantification of data from BR1008c FBLN4 IF / AZDye555-BLMP6 binding in the four origins of cores. The mean intensity of the red and green signal was scored in arbitrary units from 0 to 3 (highest). * $P<0.05$ (Anova). Error bars: SEM.

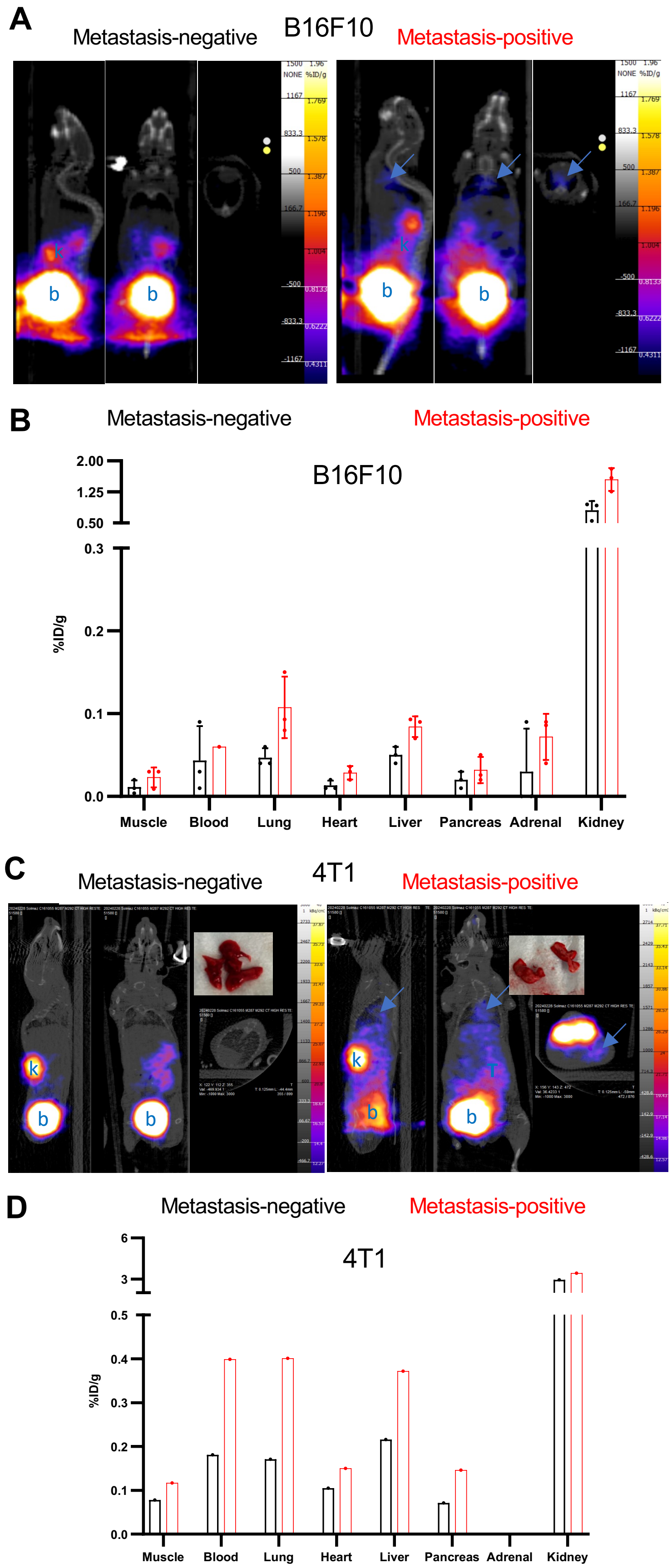


**Figure S3. A**, Radiolabeled $^{68}$Ga-BLMP6 iv-injected into C57BL/6 mice without and with B16F10 lung metastases detected by IVIS. At 1 h, PET/CT images are scaled equally. **B**, Biodistribution in tissues resected from mice in (A) with gamma counter plotted as the percentage of the injected dose per gram tissue (%ID/g tissue). **C**, Radiolabeled $^{68}$Ga-BLMP6 iv-injected into BALB/c mice without and with 4T1 lung metastases detected by IVIS. At 1 h, PET/CT images are scaled equally. **D**, Biodistribution in tissues resected from mice in (C) with gamma counter plotted as the percentage of the injected dose per gram tissue (%ID/g tissue). Arrow: $^{68}$Ga-BLMP6 in metastases. b: bladder, k: kidney.

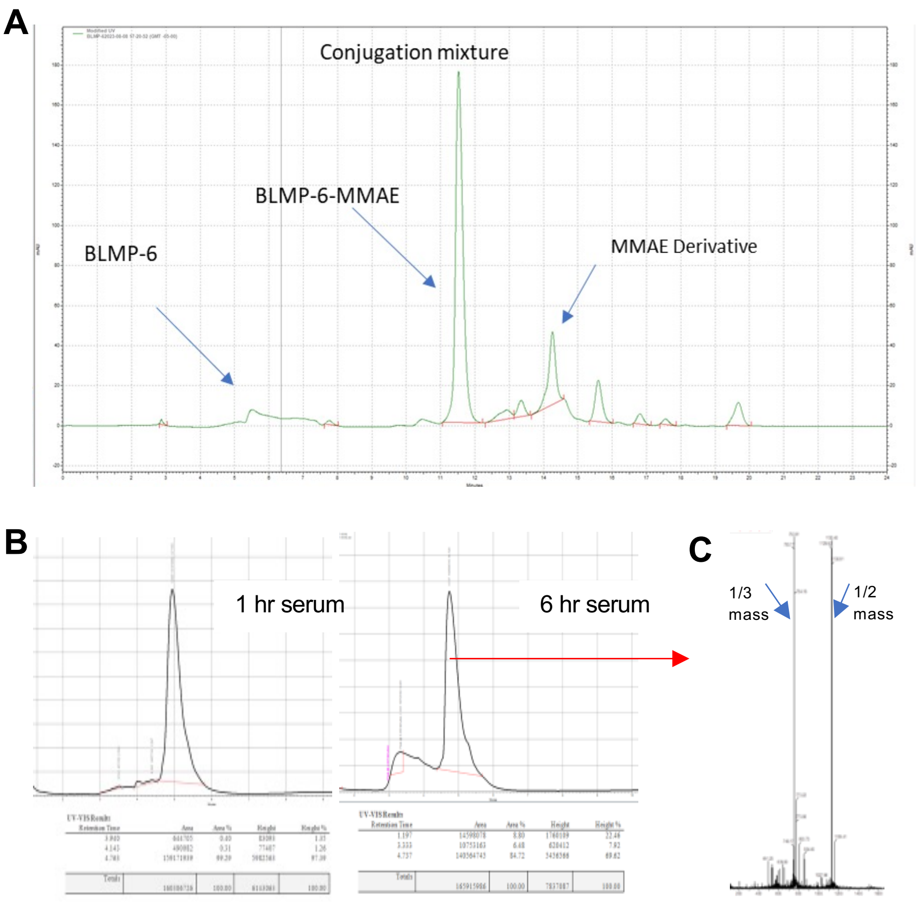


**Figure S4.**
**A**, HPLC analysis of MMAE-BLMP6 synthesis. **B**, HPLC analysis of MMAE-BLMP6 in mouse plasma after 1 and 6 h showing the main peak (taken for MS) and metabolite formation at 6 h. **C**, MS analysis confirming MMAE-BLMP6 identity *via* fragmentation patterns.
